# Psittacid Adenovirus-2 infection in the critically endangered orange-bellied parrot (*Neophema chrysogastor*): A key threatening process or an example of a host-adapted virus?

**DOI:** 10.1101/479360

**Authors:** Nian Yang, Jennifer McLelland, David J. McLelland, Judy Clarke, Lucy Woolford, Paul Eden, David N. Phalen

## Abstract

Psittacid Adenovirus-2 (PsAdv-2) was identified in captive orange-bellied parrots (*Neophema chrysogastor*) during a multifactorial cluster of mortalities at the Adelaide Zoo, South Australia, and an outbreak of *Pseudomonas aeruginosa* septicaemia at the Tasmanian Department of Primary Industries, Parks, Water and Environment captive breeding facility, Taroona, Tasmania. This was the first time that an adenovirus had been identified in orange-bellied parrots and is the first report of PsAdv-2 in Australia. To investigate the status of PsAdv-2 in the captive population of orange-bellied parrots, 102 healthy birds from five breeding facilities were examined for the presence of PsAdv-2 DNA in droppings and/or cloacal swabs using a nested polymerase chain reaction assay. Additionally, eight birds released to the wild for the 2016 breeding season were similarly tested when they were recaptured prior to migration to be held in captivity for the winter. PsAdv-2 was identified in all breeding facilities as well as the birds recaptured from the wild. Prevalence of shedding ranged from 29.7 to 76.5%, demonstrating that PsAdv-2 is endemic in the captive population of orange-bellied parrots and that wild parrots may have been exposed to the virus. PsAdv-2 DNA was detected in both cloacal swabs and faeces of the orange-bellied parrots, but testing both samples from the same birds suggested that testing faeces would be more sensitive than cloacal swabs. PsAdv-2 was not found in other psittacine species housed in nearby aviaries at the Adelaide Zoo. The source of the infection in the orange-bellied parrots remains undetermined. In this study, PsAdv-2 prevalence of shedding was higher in adult birds as compared to birds less than one year old. Preliminary data also suggested a correlation between adenovirus shedding prevalence within the breeding collection and chick survival.

## Introduction

The orange-bellied parrot (*Neophema chrysogaster*) is the most critically endangered parrot in the world [1]. It is a migratory species that breeds in south-western Tasmania and historically wintered along the coast of South Australia, Victoria, and New South Wales. It was abundant before the 1920s [2], but declined to around 200 birds in the 1990s [3]. In the spring of 2016, just 13 birds returned to the only known remaining breeding site, in Melaleuca, Tasmania [4]. While many possible causes have been suggested, the threatening processes driving this decline are incompletely understood (reviewed in Stojanovic et al. [5]). To save this species, a captive breeding program was initiated in 1984 [6]. Currently, there are over 400 birds managed in six captive breeding institutions. The wild population is augmented annually with the release of captive bred birds to the breeding ground each spring [4].

Captive breeding programs are increasingly the last resort for the survival of many endangered species [7, 8, 9]. A potential risk of captive breeding and release programs is the introduction of disease to both the captive and wild populations from other species in the captive collection or from native wild or feral species frequenting the breeding facilities [10, 11]. If an infectious disease is introduced into a captive population of endangered species it can prove challenging or even impossible to eradicate [12, 13]. The introduction of novel pathogens to wild populations can also threaten their viability [14,15,16].

Endangered psittacine birds (parrots) in captive breeding programs are particularly at risk for exposure to introduced viruses, as multiple viruses have been established in parrots as the result of the historic and ongoing trade in wild-caught parrots [17]. Notable examples of virus incursions in captive breeding programs for endangered parrots, include Avian Bornavirus in the main breeding colony of Spix macaws (*Cyanopsitta spixii*) [11] and Psittacine Beak and Feather Disease Virus (PBFDV) in captive and wild populations of Echo parrots (*Psittacula eques*) [18]. The orange-bellied parrot population is also threatened by PBFDV with three incursions of PBFDV occurring in the captive or wild populations in the past 15 years [19].

Adenoviruses also have the potential to impact captive breeding programs of endangered parrots. Adenovirus infections have been described and variably characterised in many species of parrots originating from all of their geographic distributions [20, 21, 22, 23, 24]. They have been associated with a range of lesions, including hepatitis, splenitis, pancreatitis, enteritis, nephritis, conjunctivitis and pneumonia [21, 25, 26, 27]. At least one adenovirus has been shown to cause significant ongoing mortality events in captive-raised *Poicephalis* species in South Africa, and *Poicephalis* and other species in Europe [28, 29]. In contrast, many reports describe adenovirus-associated disease where only one or a few birds are affected [24, 25, 30]. In some of these reports, adenovirus-associated lesions were part of a multifactorial disease complex [20, 21, 31, 32], or were incidental findings [29, 33, 34]. This suggests that, in at least some instances, adenovirus-associated lesions in parrots reflect reactivation of subclinical infections in the face of immune suppression, a phenomenon recognised in other species, including chickens [35].

Overall, the epizootiology of adenoviruses that affect parrots is poorly understood. It is likely, however, based on the behaviour of adenoviruses in poultry, that the adenoviruses that affect parrots are maintained in subclinically infected individuals, and that disease may only occur when birds are stressed or co-infected with immunosuppressive viruses [36, 37, 38]. It is also possible that given that many species of parrots from multiple geographic origins are commonly housed together, that some species are more likely to be subclinically infected while other species are more prone to infection resulting in disease. Evidence for this was provided in a recent report where it was shown that a novel adenovirus shed by subclinically-infected purple-crowned lorikeets (*Glossopsitta porphyrocephala*) was found to be fatal to in-contact red-bellied parrots (*Poicephalus rufiventris*) [24]. Evidence for widespread subclinical infection of parrots by Psittacid Adenovirus 2 (PsADv-2) in parrots was also demonstrated by a study of avicultural birds in Slovenia [34] where PsAdv-2 DNA was found in cloacal swabs of 10.2% (13/128 birds) birds tested, yet there was no history of adenovirus disease in these collections.

In the current study, we report, for the first time, the presence of PsAdv-2 in Australia. We demonstrate that it is widespread in the captive breeding population of orange-bellied parrots and that the wild population has been exposed. We also provide preliminary data indicating that subclinical infection may result in reduced chick survivability and that PsAdv-2 disease is most likely to occur in birds with other concurrent diseases.

## Materials and methods

### Animal ethics

All the material used in this study was submitted for diagnostic purposes. The Animal Ethics Committee at the University of Sydney was informed that findings from the diagnostic material were to be used in a publication and a formal waiver of ethics approval has been granted.

### Investigation of mortality events at two captive breeding facilities

The first mortality event (n = 8) occurred between February- and March 2016 in the captive breeding collection at the Adelaide Zoo, South Australia (34°54’46.71“S, 138°36’25.00”E). The second mortality event (n=25) occurred in January 2017 at the Department of Primary Industries, Parks, Water and Environment orange-bellied parrot breeding facility in Taroona, Tasmania (42°57’0.70“S, 147°21’13.71”E) [39]. Representative tissues from deceased birds were collected into 10% neutral buffered formalin, embedded in paraffin, sectioned at 4 µm and stained with haematoxylin and eosin. Liver, kidney and/or spleen were frozen at −20°C from all eight birds from the Adelaide mortality event, and15 of the 25 birds that died at the Taroona facility.

### Detection of adenovirus DNA in tissues

Tissues from two birds from the Adelaide Zoo and 15 birds from the Taroona breeding facility were submitted for PCR screening for adenovirus infection (Table 1). DNA was extracted from all tissues submitted with a DNA extraction kit (DNeasy Blood & Tissue Kit, Qiagen, Doncaster, Victoria, Australia) following the manufacture’s recommendations. Adenovirus DNA was detected using nested-PCR with degenerated primer sets described by Wellehan et al. [40]. The first and nested rounds of the DNA amplifications were based on the following protocol: One cycle of 95 C∘ for 3 min, followed by six cycles where the first annealing temperature (60°C) was decreased by 2°C with each cycle. In all subsequent cycles (n=44) the annealing temperature was 50°C. Primers were allowed to anneal in all cycles for 30 s and all extension phases were at 72°C for 30 s., these were followed by a denaturation phase at 95°C for 30 s. Following the last standard cycle, the samples were held at 72°C for 2 min and immediately chilled to 5°C. The amplification products were separated by electrophoresis on 1.5% agarose gels containing ethidium bromide and visualized under ultraviolet light. The second amplification products of the positive samples were purified by centrifugation (Amicon Ultra Centrifugal Filtration Units, Millipore, Tulagreen, Ireland) and sequenced in both directions using the amplification primers (Australian Genome Research Facility, Sydney, New South Wales, Australia). Sequences were compared with adenovirus sequences in GenBank using NCBI BLAST [41].

**Table 1.**
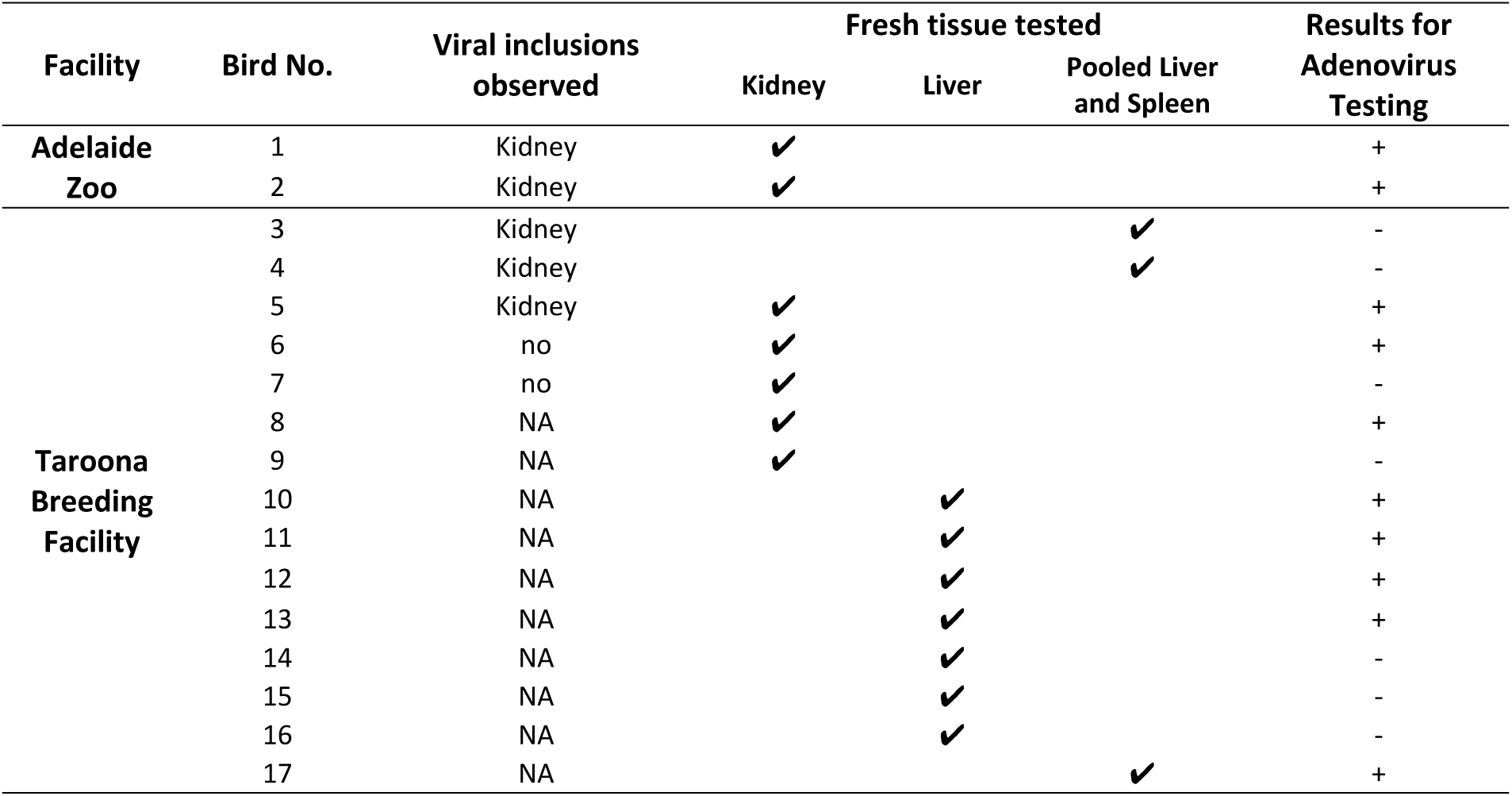
Histopathological findings and PCR results of orange-bellied parrots (*Neophema chrysogaster*) necropsied at Adelaide Zoo and Taroona Breeding Facility.

### Detection of adenovirus DNA in cloacal swabs and droppings

Cloacal swabs, frozen dry at −20°C prior to analysis, were obtained from 102 outwardly healthy captive orange-bellied parrots at Adelaide Zoo (n = 23), Priam Parrot Breeding Centre (n = 17) (35°20’22.71“S, 149°15’0.65”E), Healesville Sanctuary (n = 37) (37°40’57.13“S, 145°31’51.20”E), and Moonlit Sanctuary (n = 25) 38°12’41.70“S, 145°15’2.89”E). This sampling represented 92% of the Adelaide Zoo flock, 100% of the Priam flock, and approximately 50% of the Healesville and Moonlit flocks. Faecal samples were collected concurrently from 35 birds at the Healesville Sanctuary. Eight captive-raised female birds released to the wild for the 2016-2017 breeding season were recaptured prior to migration and held in isolation over winter in captivity at Werribee Open Range Zoo (37°55’22.23“S, 144°40’2.58”E). Cloacal swabs were collected from all eight birds.

An additional 38 cloacal swabs from adjacently housed parrots at the Adelaide Zoo were also tested. Species tested included: elegant parrot (*Neophema elegans*) (n=15); regent parrot (*Polytelis anthopeplus*) (n=4); black-capped lory (*Lorius lory*) (n=4); eclectus parrot (*Eclectus roratus*) (n=5); scarlet macaw (*Ara macao*) (n=2); crimson-bellied parakeet (*Pyrrhura perlata*) (n=6); blue-and-yellow macaw (*Ara ararauna*) (n=1) and yellow-crowned amazon (*Amazona ochrocephala*) (n=1).

DNA extraction, PCR amplification of adenovirus DNA, and sequencing were done as described above. To test the hypothesis that the amplification products were from PsADv-2, PsAdv-2 specific primers (3’GAACAGAGAGGAGGAAGG, 5’GGGAAAACCGAAAAAGAGCA) were designed to amplify the second amplification products of the positive samples using Geneious (Biomaters, Auckland New Zealand). One positive sample from each tested institution was randomly selected for sequencing with the PsAdv-2 specific internal primers (Australian Genome Research Facility, West Mead, Australia).

### Characterisation of suspected adenoviruses in the black-capped lories

The PCR amplification products of the cloacal swabs from 3 black-capped lories were of the appropriate mass to be adenovirus. The attempt to sequence the amplicons with the degenerate amplification primers failed. To test the hypothesis that the amplification products were from PsADv-2, two additional PsAdv-2 specific primers were designed (3’ACAGAGAGGAGGAAGGCAGT, 5’TCACACACCCTTTGCCCTTT) using Geneious and amplification was attempted using amplicons from the first PCR reaction for the three lories and amplicons from the orange-bellied parrots as positive controls.

### Sensitivity of the PCR assay

To determine the sensitivity of the PCR assay, the first amplification products were purified. Then 10-fold serial dilutions of the purified product were made, starting from 10^5^ copies/μL to 1 copy/μL. The dilutions were then amplified by the nested PCR assay.

### Data analysis

Sex, age and reproductive data (n= 65 birds) for the 2016-17 breeding season of the birds were acquired from the breeding institutions. The age of birds was categorised as adult (≥ 1 yr) and juvenile (<1yr) groups. Significant tests for associations between the presence of the adenovirus infection and the age and sexes of the tested birds were based on Chi-squared test (p ≤ 0.05).

Linear regression modelling [42] was used to assess the correlations between the prevalence of adenovirus and indicators (fertility of eggs, egg hatchability, percentage of chicks that fledged) of reproductive success among four of five breeding collections (Healesville Sanctuary, Adelaide Zoo, Moonlit Sanctuary and Priam Parrot Breeding Centre). Goodness of fit was assessed by the R^2^ values. Statistical significance of the linear regression coefficients was assessed by the P-value from the *t-test*.

## Results

### Investigation of mortality events at two captive breeding facilities

#### Adelaide Zoo

Birds were either found dead, or died after a short period of hospitalization, having been found weak, some with increased respiratory effort. The only consistent gross necropsy finding was thin body condition and birds were underweight, with empty gastrointestinal tracts. The first bird to die in February 2016 was diagnosed with systemic Gram-negative bacterial infection. Intra-nuclear inclusion bodies were observed in hepatocytes and were associated with mild hepatocellular necrosis. Electron microscopy identified viral particles 90-100nm in diameter that were morphologically consistent with an adenovirus. Four birds had mild non-specific histopathological changes and one bird had necrotizing Gram-negative sinusitis. Locally abundant intra-nuclear inclusions in renal epithelial cells of the collecting ducts associated with mild to moderate degenerative changes were found in two birds (Fig 1) that had severe aspergillosis; Both of the birds were positive for adenovirus DNA by PCR (Table 1). The mortality event was attributed to cumulative stressors in the flock, including a period of hot and unusually humid weather. Adenoviral lesions in these cases were considered secondary to immunosuppression in debilitated birds, and were not considered to have contributed significantly to the deaths of these birds.

**Fig 1.**
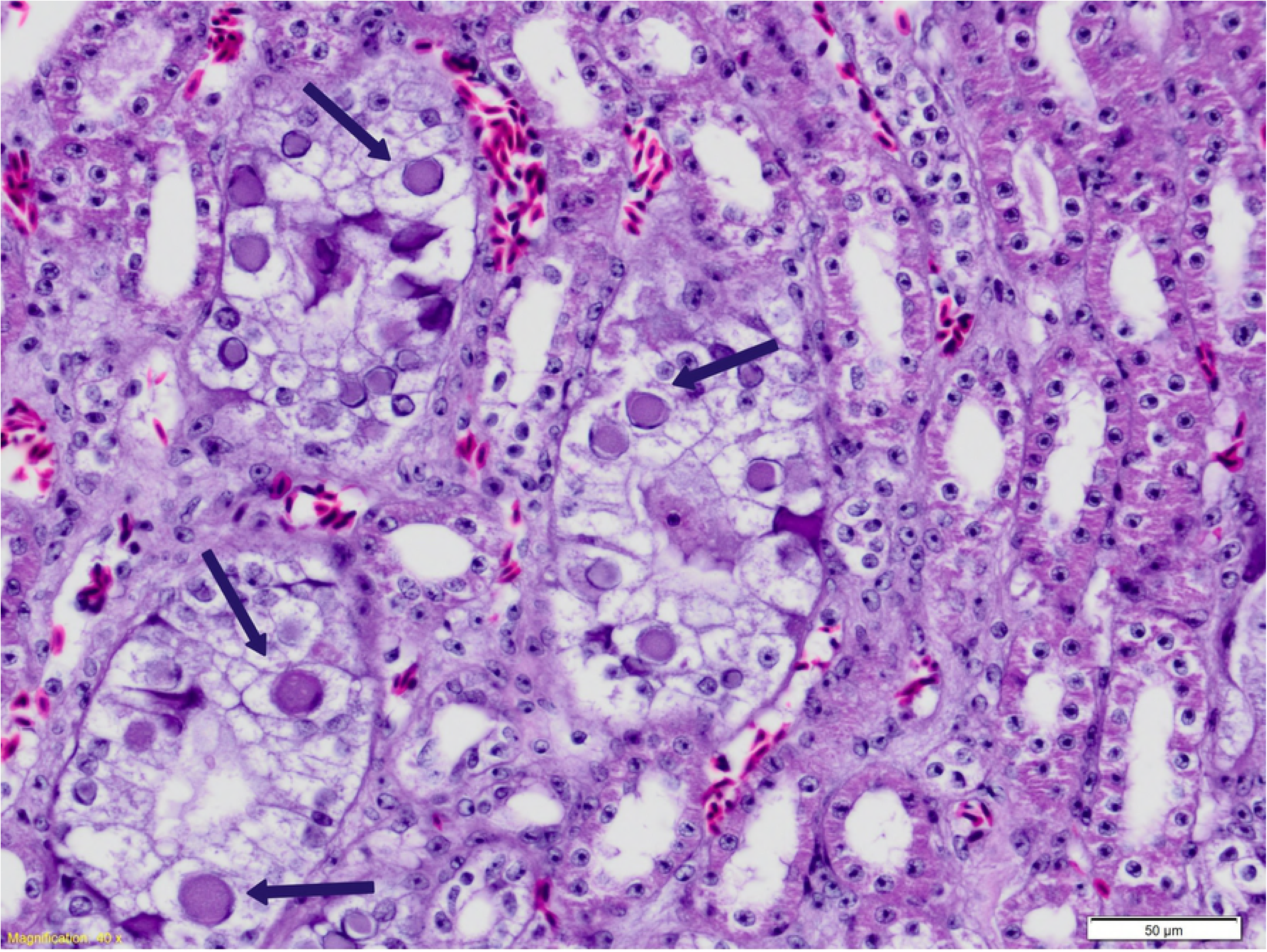
Intra-nuclear inclusion bodies in renal collecting duct epithelial cells in an orange-bellied parrot infected with psittacid adenovirus 2 (Arrows).

#### Taroona breeding facility

The mortality event in Taroona was attributed to *Pseudomonas aeruginosa* septicaemia traced to contaminated sprouted seed [39]. Histopathology was available for five birds; intra-nuclear inclusions were found in kidney sections of three birds. Eight of the fifteen birds were PCR positive for PsAdv-2, with virus identified in kidney and/or liver. Two birds with renal inclusions tested PCR negative (pooled liver and spleen); one PCR-positive bird (liver) had no viral inclusions identified (Table 1).

#### Sequencing and PCR amplification with PsAdv-2 specific primers

The sequences (323bp) of the amplicons from the orange-bellied parrots at the Adelaide, Taroona, breeding facilities were all identical to the sequence of PsAdv-2.

#### Detection of adenovirus DNA in cloacal swabs and droppings

Adenovirus infection was detected in all institutions, including the recaptured birds held at Werribee Zoo. The overall prevalence of the adenovirus infection was 42.7%, and the prevalence for each institution ranged from 29.7% to 76.5% (Table 2). The sequencing results of the DNA samples from each institution were all identical to PsAdv-2. All amplification products from positive orange-bellied parrots re-amplified with nested specific PsADV-2 primers. Three of four black-capped lories were PCR-positive for adenovirus; the other 35 parrots at Adelaide Zoo were negative. The sequences of the amplification products from the black-capped lories were not readable and amplification products were not generated using PsAdv-2 specific primers.

**Table 2.**
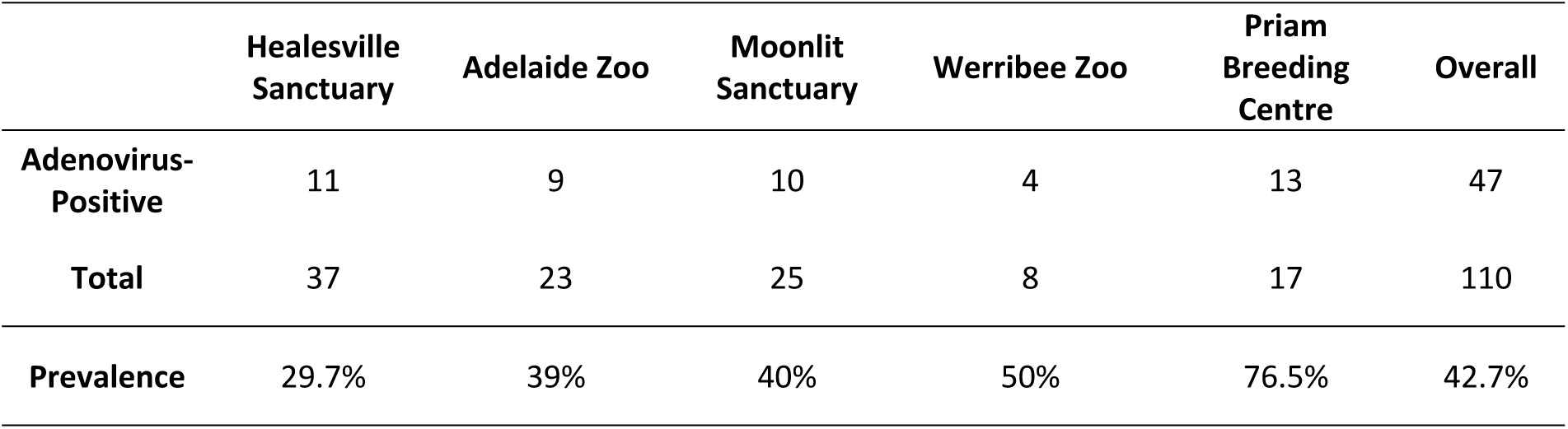
Detection of psittacid adenovirus 2 in cloacal swabs from orange-bellied parrot (*Neophema chrysogaster*) across five captive breeding institutions.

|  | Healesville<br>Sanctuary | Adelaide Zoo | Moonlit<br>Sanctuary | Werribee Zoo | Priam<br>Breeding<br>Centre | Overall |
| --- | --- | --- | --- | --- | --- | --- |
| **Adenovirus-**<br>**Positive** | 11 | 9 | 10 | 4 | 13 | 47 |
| **Total** | 37 | 23 | 25 | 8 | 17 | 110 |
| **Prevalence** | 29.7% | 39% | 40% | 50% | 76.5% | 42.7% |

#### Relative sensitivity of sample sources and sensitivity of PCR assay

A comparison of PCR results for the paired faecal samples and cloacal swabs collected from 35 orange-bellied parrots at Healesville Sanctuary is shown (Table 3). Seven birds were both positive on both samples; 25 birds were negative on both samples. Three birds were only positive for adenovirus in faeces.

**Table 3.**
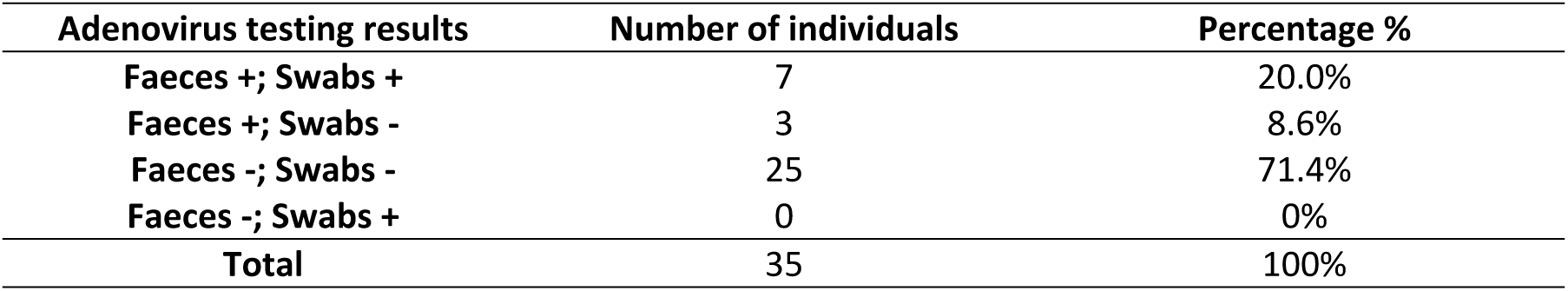
Relative sensitivity of faeces and cloacal swabs for the detection of adenovirus DNA in orange-bellied parrots (*Neophema chrysogaster*).

#### Sensitivity of the nested PCR assay

All dilutions of DNA could be detected by the nested PCR indicating that it was able to detect a minimum of six copies of PsAdv-2.

#### Correlation between infection prevalence and age, sex and reproductive data

The presence of PsAdv-2 was significantly associated with age, with adult birds more likely to be infected by PsAdv-2 than birds < 1 yr 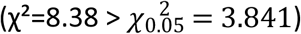. There was no significant association between PsAdv-2 infection and sex 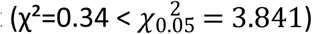. A quasi-significant negative correlation was observed between the fledgling rate of hatched chicks and the PsAdv-2 prevalence at the institutional level (*R*^2^ = 0.890, *p_α_* = 0.057, *p_β_* = 0.013) (Fig 2). No significant association was observed between PsAdv-2 prevalence and fertility rate or hatch rate of fertile eggs.

**Fig 2.**
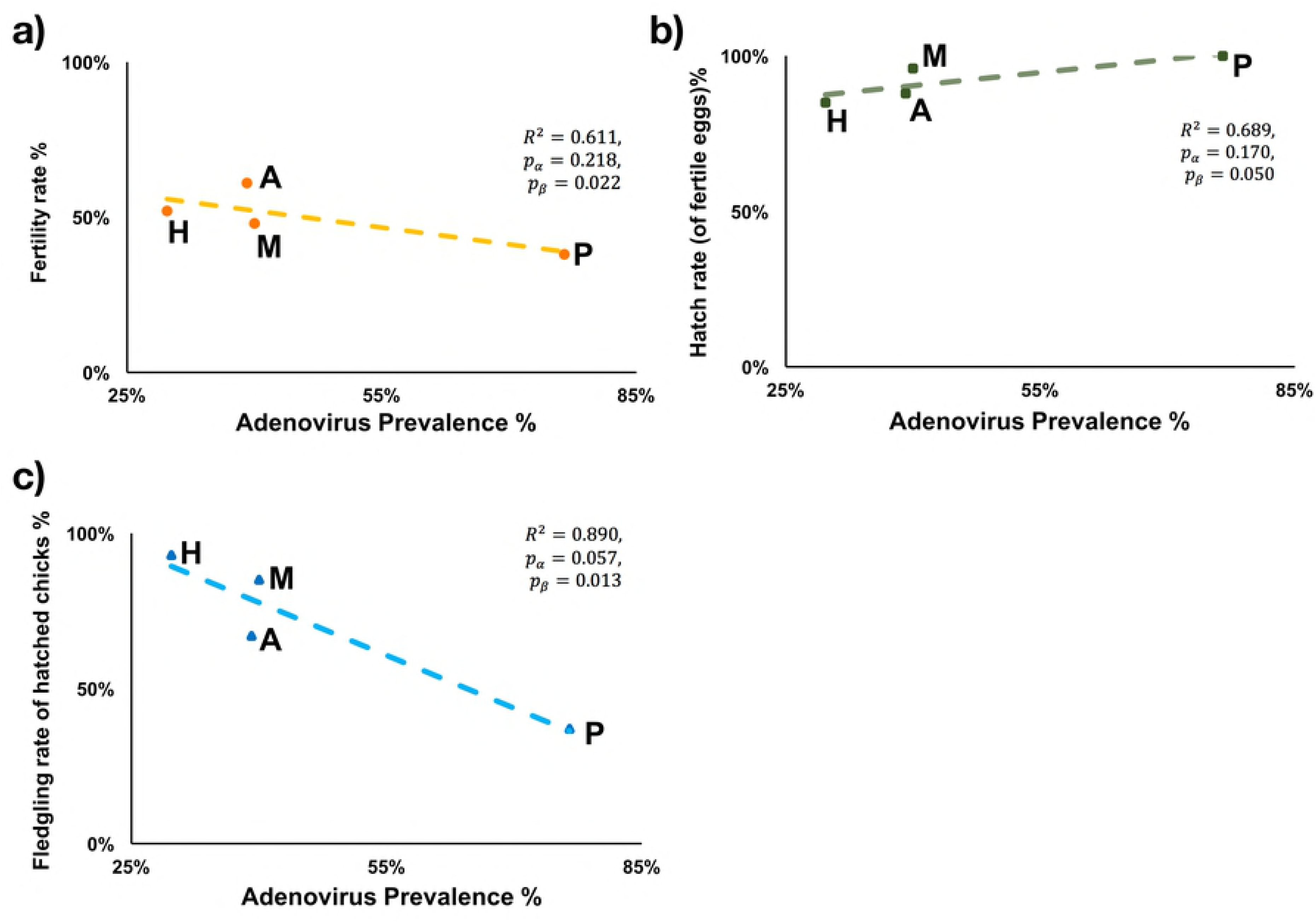
Linear regression model of the correlations between adenovirus prevalence and reproductive success of the orange-bellied parrots (*Neophema chrysogaster*) during the 2016-17 breeding season. (H: Healesville Sanctuary, A: Adelaide Zoo, M: Moonlit Sanctuary, P: Priam Parrot Breeding Centre).

## Discussion

The orange-bellied parrot is on the verge of extinction in the wild. Efforts to maintain a wild population require the annual release of captive-raised birds [4]. Detailed protocols are in place, including the isolation of breeding stocks from other captive parrots, testing and quarantine, to minimise the impact of diseases in the captive breeding program and to prevent the introduction of disease from the captive-raised orange-bellied parrots to the wild orange-bellied parrots [43]. The identification of adenovirus inclusions in birds dying from a multifactorial disease at the Adelaide Zoo, and from *Pseudomonas areuginosa* septicaemia at the Taroona breeding facility prompted efforts to determine the specific adenovirus that was causing these lesions, determine how widespread infection was in the captive breeding stock, and if the wild birds had already been exposed.

Sequencing data generated in this study shows that the cause of the lesions in the Taroona and Adelaide birds, and the adenovirus subclinically infecting other orange-bellied parrots, is the Siadenovirus PsAdv-2. This is the first record of this virus in Australia and the most comprehensive investigation into its epizootiology to date. PsAdv-2 was first detected in North America in a plum-headed parrot (*Psittacula cyanocephala*) that died with an acute systemic bacterial infection [21]. This bird also had adenovirus inclusions in the liver that were associated with a mild inflammatory response and individual hepatocyte necrosis. Wellehan et al. [21] also reported PsAdv-2 infection in an umbrella cockatoo (*Cacatua alba*) that died with a chronic encephalitis of unknown cause. Adenovirus inclusions were not seen in this bird and there was no evidence that PsAdv-2 contributed to its illness. The only other study of PsADV-2 involved a survey of 128 apparently healthy captive parrots in Slovenia [34]. In this study, cloacal swabs from 13 (10.2%) of the parrots were positive for PsADV-2 DNA. The positive species originated from South America, Africa, Southwest Asia, and Australia, including a *Neophema* sp. [34].

Based on the study in Europe [34] and our findings in orange-bellied parrots, PsAdv-2 has a wide host range and is able to circulate in parrot populations without causing apparent disease, or doing so infrequently. While it will be challenging to determine the species of parrots from which this virus originated, it appears that its ability to cause subclinical infections in a wide range of parrot species has resulted in its global distribution [21, 34]. Finding virus in cloacal swabs and droppings suggests that faecal/urinary-oral transmission would be the most likely form of transmission. However, some adenoviruses can be transmitted vertically [44], so it is possible that PsAdv-2 may also be transmitted this way.

Whether PsAdv-2 was introduced into the orange-bellied parrot population by indirect contact with other native or exotic species, or whether it originated in the orange-bellied parrot, is not known. All institutions housing orange-bellied parrot breeding populations, with the exception of the Taroona facility, have multiple other species of both native and exotic birds, and many of the enclosures are open to the environment and thus have the potential for wild bird exposure. We were only able to test parrot species in nearby enclosures at one institution and we found no evidence of infection to suggest they were the source of infection or that they had been infected with PsAdv-2 from the orange-bellied parrots. More extensive testing of other birds housed adjacent to the orange-bellied parrots at the other breeding institutions is warranted to further investigate potential transmission pathways.

Amplicons of the appropriate mass were detected in three of four black-capped lories, but attempts to sequence these products were unsuccessful. These amplification products are not PsAdv-2 DNA as PsAv-2 specific primers did not produce an amplicon when used in a nested reaction with the first amplification product. Therefore, they may represent non-specific amplification products or the adenoviruses that infected the black-capped Lories were not PsAdv-2. Cloning and sequencing the amplification products will be necessary to differentiate between these two possibilities.

While most PsAdv-2 infections appear to be subclinical, they may not be completely innocuous. The first report of this infection, in a plum-headed parrot [21], described a mild chronic hepatitis associated with inclusions and we saw a similar lesion in one of the orange-bellied parrots that died with aspergillosis in the Adelaide Zoo mortality cluster. Renal inclusions were also seen in two other birds from the Adelaide Zoo and three of five birds for which there was histopathology that died from *Psuedomonas auregenosia* had renal inclusions. Virus inclusions in the kidneys varied between being uncommon to locally abundant and while generally not associated with necrosis nor a significant inflammatory response, degeneration of tubular epithelial cells was seen. This variation in lesions suggests that while adenovirus infection may persist in one or more organs in healthy birds, virus replication is triggered or increased in birds that are compromised by other infectious diseases and at these times it may have an impact on the host. Given that the immunosuppressive PBFDV is enzootic in the captive orange-bellied parrot breeding population [19], future studies correlating PsAdv-2 shedding with past and current PBFDV infection status would be warranted.

The reproductive success of the captive breeding program of orange-bellied parrots is relatively poor with problems including infertile eggs and suboptimal chick survival after hatch [5]. This may reflect a lack of genetic diversity in the overall population, challenges associated with breeding orange-bellied parrots outside of their normal summer range, malnutrition and/or other unrecognized factors. In this paper, we present preliminary data suggesting a correlation between adenovirus shedding prevalence with in a breeding facility and chick survival. Additional studies correlating the infection status of the parents with chick infection status and survival will be required before the impact of PsAdv-2 on chick mortality can be determined. Testing dead, in shell, embryos and infertile eggs for the presence of PsAdv-2 would also prove useful, as it could be used to determine if egg transmission occurs and if egg infection impacts embryo survival. If egg transmission does occur, it would make eradication of the virus from the orange-bellied parrots challenging or even impossible.

A Veterinary Technical Reference Group containing members from all the institutions involved in the orange-bellied parrot captive breeding program has been established to provide advice to the Orange-bellied Parrot Recovery Team. The value of this Reference Group was demonstrated in their response to the initial detection of an adenovirus in the captive orange-bellied parrots. The Reference Group was instrumental in making samples available for the adenovirus testing done in this study. Additionally, the Reference Group agreed prior to testing that if an adenovirus was found to be present in all institutions and wild birds had been exposed, then it would be considered to be endemic in both the captive and wild populations and no control efforts would be undertaken. Our findings show that PsAdv-2 is indeed endemic in the captive orange-bellied parrot population and that the wild population has been exposed. Our findings of an overall prevalence of 42.7% infection in the tested birds also make any type of control option virtually impossible. Isolation of nearly half of the birds and taking this number of captive birds out of the breeding population would not be practical and likely would result in a loss of genetically important breeding stock. Also, the sensitivity of PCR testing of cloacal swabs or droppings to detect infected birds is unknown. It is possible that infected birds shed virus intermittently or may not shed virus at all. We also found that virus shedding was more common in adult birds than birds less than one year. This may mean, as occurs in adenovirus infections in poultry, that although birds are infected with an adenovirus, shedding is more likely to occur in sexually mature birds (McFerran and Smyth, 2000). If any of these hypotheses are correct, the prevalence of infected birds may be much higher than the results of this study have documented. Repeated testing of birds over time, and opportunistic parallel testing of cloacal swabs/droppings and tissues at necropsy, will be necessary if the sensitivity of this test is to be determined.

This study generated additional results of potential diagnostic significance. While both cloacal swabs and droppings were found to be useful samples for detecting PsAdv-2 infection in the orange-bellied parrot, in the 35 samples where both cloacal swabs and droppings were submitted, 20% of the cloacal swabs were positive, while 28.7% of the droppings were positive. This suggests that droppings may be the preferred sample to test when screening for PsAdv-2 infection, eliminating the need to catch a bird to test it. Additional diagnostic findings show that both the liver and the kidney can contain PsAdv-2 DNA without the presence of inclusion bodies in histological sections. Also, PsAdv-2 DNA can be found in some, but not all, liver samples, from birds with renal inclusions, making kidney the organ of choice when testing cadavers for the presence of PsAdv-2.

It has been proposed that private aviculturists might be used to assist in breeding the orange-bellied parrot to increase the numbers in captivity and the availability of birds for release into the wild. The findings of this study argue against this proposal. Given the uncertainty of the sensitivity of testing for PsAdv-2, it would be impossible to guarantee that birds sent to aviculture collections would be free of this virus and therefore could pose a risk to the aviculturists’ birds. Likewise, while extensive testing is done yearly for the PBFDV in the captive orange-bellied parrot population [19], it has been impossible to eradicate it and so these birds remain potential sources of this virus. Introduction of new infectious diseases into the captive orange-bellied parrot population in aviculturists’ collections is also a risk. Most aviculturists have limited resources and it would be unlikely that they could provide housing and management practices that would prevent disease transmission from their own birds to the orange-bellied parrots. This is of great concern in that Avian Bornavirus-2 and 4 and Psittacid Herpesvirus-3, pathogens that have had major impacts on avicultural species globally, are known to be present in some Australian avicultural collections [45, 46). Also, new viruses are still being discovered in Australian aviculture collections and the epizootiology of these viruses and means of testing for them are largely unknown [24, 47].

In conclusion, this study documents the presence of PsAdv-2 in Australia for the first time. It also shows that it is endemic in the captive orange-bellied parrot population and that the wild population has been exposed. PsAdv-2 does not appear to be highly pathogenic to the orange-bellied parrot, but reactivation of replication may occur in birds that are experiencing other stressors or concurrent infections. Preliminary findings hint that PsAdv-2 may impact chick survival to fledging, but a more detailed investigation will be required to confirm this. Because PsAdv-2 is endemic in the orange-bellied parrot population, its impact on the population is minimal or unproven, and the sensitivity of available testing methods is not known, efforts to control this virus in this population are unlikely to be successful and are not recommended at this time.

## Acknowledgements

This research was funded by the Sydney School of Veterinary Sciences, The University of Sydney. The authors are grateful to the staff of Adelaide Zoo, Taroona Orange-bellied Parrot Breeding Facility, Healesville Sanctuary, Moonlit Sanctuary, Priam Parrot Breeding Centre and the Werribee Open Range Zoo who collected samples for this study. The authors thank Annika Everaardt, Lisa Tuthill, Leanne Wicker, and Daniel Gowland for their support and providing samples.

